# An investigation of the IL-23/Th17 axis and transcriptomic profiles of T helper subsets in endometriosis

**DOI:** 10.64898/2026.06.11.731688

**Authors:** Danielle J. Sisnett, Katherine B. Zutautas, Jaelis P. Holmes, Jessica Pudwell, Kira King, Olga Bougie, Chandrakant Tayade

## Abstract

Endometriosis (EMS) is a chronic inflammatory disease characterized by ectopic endometrial-like tissue growth and immune dysregulation. Aberrant T cell responses are implicated in EMS pathogenesis, however, functional reprogramming of T helper (Th) subsets across disease stages remain unclear. We profiled systemic and local immune mediators (cytokines/chemokines) and performed bulk RNA sequencing to comprehensively characterize Th1, Th1/17, and Th17 cell subsets from EMS patients and healthy controls. Across patient plasma, peritoneal fluid (PF), and matched eutopic and ectopic tissues, we observed systemic and local cytokine/chemokine alterations, including elevated IL-6 (plasma), FLT-3L and G-CSF (eutopic), and IL-1RA and IL-23 (p40; PF) in severe-stage EMS. Flow cytometry depicted elevated pathogenic Th17 cells in patient PF compared to matched non-pathogenic Th17 and Treg cell subsets. Additionally, circulating Th17 cells were increased in patients with mild (stages I-II) relative to severe (stages III-IV) EMS. RNA sequencing revealed extensive Th subset reprogramming in EMS, most predominately in Th17 cells (2,220 DEGs). Collectively, we reveal significant immune remodeling in EMS and highlight a distinct, aberrant Th17 cell phenotype. Results support repurposing of IL-23- and IL-17-targeted therapeutics, already effectively implemented in other chronic inflammatory diseases, to broaden therapeutic options for EMS and associated comorbidities.

## Introduction

Endometriosis (EMS) is an estrogen-dependent, chronic inflammatory disease, wherein endometrial-like tissue grows ectopically, resulting in inflammation, fibrosis, and scar tissue. Impacting approximately 190 million reproductive-age women worldwide, and an undisclosed number of gender-diverse individuals, this disease holds a significant social and economic burden^1^. EMS commonly presents clinically as chronic pelvic pain, dysmenorrhea, and infertility, with up to 50% of global cases of female infertility attributed to EMS^1,2^. However, due to symptom heterogeneity, invasive diagnostic measures, and the stigmatization of reproductive disorders, diagnosis is often delayed up to 12 years, exacerbating disease burden^3^. Moreover, as therapeutics are still quite limited, either being invasive, short-lasting, and/or hormonal in nature, there is a great need to expand therapeutic options for patients.

EMS etiology and pathogenesis are still not completely understood, likely due to the complex interplay between hormones, genetics, and the immune system. Immune dysregulation is commonly reported in patients and recognized to enable key hallmark features of disease, such as inflammation, angiogenesis, and proliferation, ultimately facilitating lesion survival and growth. Particularly, dysfunctional T cell subsets are thought to play a dominant role in the lesion microenvironment. While the individual involvement and interplay between T helper (Th) subsets in EMS pathogenesis has not been completely elucidated, the role of Th17, Th1, and T regulatory (Treg) cell subsets have been highlighted in various studies. Indeed, we, and others, report a dysregulated IL-23/Th17/IL-17 axis in EMS, with implications to disease pathophysiology^4–9^. Specifically, IL-23 was significantly increased in patient plasma and known to promote a proinflammatory, IL-17-producing, Th17 phenotype known as pathogenic Th17 cells (pTh17)^5^. In contrast, Th17 cells not under the influence of IL-23 are more anti-inflammatory, producing IL-10, and known as non-pathogenic Th17 cells (npTh17)^10,11^. It is important to note that pTh17 cells can also co-produce IFNγ, promoting a more proinflammatory Th1 response. Due to this, this subset is commonly referred to as Th1/17 cells^12,13^. While data supports increased Th17 cells and IL-17 in EMS^6,7,14^, which is associated with disease severity, no studies have yet teased out the specific roles of pTh17 and npTh17 cells in EMS.

Under healthy conditions, Tregs work to maintain immune homeostasis, opposing proinflammatory signals from Th17 cells. However, an imbalanced ratio of Th17/Treg cells is associated with EMS^5,15,16^, likely resulting in immune dysfunction. While still largely unclear, it is proposed that an initial Th1/Th17-dominant proinflammatory microenvironment allows for early lesion establishment, which then transitions into an anti-inflammatory, Th2/Treg-dominant microenvironment to facilitate lesion growth and survival^17–19^. However, due to the transient nature of these T cell subsets, Th17 cells for example may differentiate into Tregs and vice versa dependent on environmental cues^20,21^, adding complexity to this immune landscape.

Ultimately, previous studies have focused mostly on assessing T cell frequency and cytokine/chemokine levels in EMS. Due to this, there is very limited data on transcriptomic-level changes in purified T cell populations in EMS. Thus, it is still largely unclear how key T cell subsets, including Th1, Th1/17, and Th17 cells, are reprogrammed in EMS and contribute to disease pathophysiology. In this study, we aimed to comprehensively characterize cytokines/chemokines locally (peritoneal fluid [PF], tissues) and systemically (plasma) in EMS patients compared to healthy controls and identify differentially expressed genes (DEGs) and pathways that may contribute to T cell dysfunction in EMS.

## Methods

### Human subjects

EMS patients (n=22) were recruited from gynecology clinics at the Kingston Health Sciences Center (KHSC, Canada). To meet eligibility criteria, participants were between 30-50 years old, had a uterus, at least one ovary, and had excision surgery scheduled within this study period. All individuals in the EMS cohort had a pre-existing diagnosis of EMS prior to study enrollment based on previous surgical findings, imaging (ultrasound and/or MRI), clinical history/symptoms, or a combination of these three criteria (**Table S1**). Peripheral blood was collected prior to surgery. During surgery, PF and endometrial biopsies of ectopic and eutopic tissues were collected. Histopathological analysis of excised lesions was subsequently performed to confirm EMS diagnosis. At this time, surgical staging was conducted by the main surgeon using the revised American Society of Reproductive Medicine (rASRM) staging system^22^. While this system is criticized due to poor correlation of rASRM staging criteria and meaningful clinical outcomes^23^, this is still the most widely accepted classification system^24^.

Healthy control participants (n=19) were enrolled using community postings, online advertisement, and word of mouth. These individuals had no history of EMS or other gynecological conditions, such as fibroids, pelvic inflammatory disease, or chronic pelvic pain. Individuals with cardiovascular disease at baseline, diabetes/metabolic syndrome, Raynaud’s phenomenon, collagen-vascular disease, or those using statins were excluded from both the EMS and control cohorts. Participants were also screened for other relevant factors, such as hormonal use and history of pregnancy, using a self-questionnaire (**Table S1**).

### Sex as a biological variable

EMS is an estrogen-dependent disease that predominately affects individuals of female sex. Accordingly, sex was considered as a biological variable in this study, with all human study participants and EMS patients included being of female sex, as assigned at birth. Participants were limited to cisgender women due to recruitment constraints. Thus, gender- and sex-diverse individuals (e.g. transgender, non-binary, or intersex people, including those undergoing gender-affirming surgery or receiving gender-affirming hormonal therapies) were not represented in this study. We acknowledge this limitation and emphasize the importance of promoting diversity and inclusivity in health research to generate data relevant to diverse global populations.

### Plasma, PBMC isolation, and multiplex cytokine array

Whole blood (18mL) was collected from participants into BD Vacutainer® tubes (CABDL366643L, VWR, USA). Blood was then diluted (1:1) with 2% FACS buffer (PBS with 2% FBS; 97068-091, VWR, Canada) and layered on top of Lymphoprep™ (07811, StemCell Technologies, Canada) within SepMate™-50 PBMC Isolation Tubes (85460, StemCell Technologies, Canada). Samples were centrifuged for 20mins at 1,200RCF at 18°C, as per manufacturer’s guidelines. Plasma was aspirated and stored at −80°C prior to multiplex cytokine array (HD48-plex, EveTechnologies, Canada), detecting 48 key immunological analytes. PBMCs were collected and washed with 2% FACS buffer, prior to *in vitro* stimulation or cryopreservation (80% FBS and 20% DMSO) for later FACS sorting.

### *In vitro* stimulation and expansion of PBMCs to Th17 cell fate

Fresh PBMCs were centrifuged at 400RCF for 10mins at 18°C (brake off) prior to resuspending in PBMC media consisting of RPMI-1640 (11875085, ThermoFisher, Canada) supplemented with 10% FBS, 1% penicillin and streptomycin (15140122, ThermoFisher, Canada), and 2% HEPES (15630080, ThermoFisher, Canada). Cells were seeded in 24-well plates at 0.5×10^6^ cells/well. Four treatment groups were used: i) Th17 cocktail and activation, ii) activation alone, iii) Th17 cocktail alone, and iv) media control. Briefly, activation consisted of plate-bound anti-human CD3 mAb and anti-human CD28 mAb, while the Th17 cocktail was comprised recombinant human (rh)TGF-β1, rhIL-1β, rhIL-6, rhIL-21, rhIL-23, anti-IFNγ, and anti-IL-4, as detailed in our previous publication^5^. Cells were incubated for 4 days at 37°C with 5% CO_2_. Following this, PBMCs were re-stimulated with 50ng/mL of PMA (74042, StemCell Technologies, Canada) and 250ng/mL of ionomycin (73722, StemCell Technologies, Canada), incubated for a further 5h, and then cells were collected for flow cytometry (panel 1).

### CD4+ T cell isolation, FACS sorting, and bulk RNA sequencing

Prior to sorting cells (panel 2), PBMCs were briefly thawed in a water bath at 37°C. Cell suspensions were added to warm RPMI media with 10% FBS and centrifuged at 400RCF for 10mins at 20°C prior to resuspending in 2% FACS buffer and counting cells. CD4+ T cells were then magnetically isolated using a commercially available kit (17952, StemCell, Canada), as per manufacturer’s protocol. CD4+ T cells were washed and resuspended in warm TexMACS media (130-097-196, Miltenyi Biotec, USA) prior to seeding in 96-well plates at 1×10^6^ cells/well. Cells were then re-stimulated prior to staining using 30ng/mL of PMA and 1μg/mL of ionomycin and incubated for 4h at 37°C with 5% CO_2_. Following incubation, IFNγ-secreting and IL-17-secreting cells were fluorescently labelled using IFN-γ Secretion Assay – Detection Kit (FITC) and IL-17 Secretion Assay – Detection Kit (PE) from Miltenyi Biotec (**Table S2**), as per manufacturer’s protocol. This was conducted to avoid permeabilizing cells to allow for downstream analysis. During staining, cells were also labelled with CD3 and CD4 extracellular Abs, as well as the viability dye (**Table S2**). Following washing, cells were resuspended in 2% FACS buffer for FACS sorting. Cells were sorted into viable populations of Th17 (CD3+CD4+IFNγ-IL-17+), Th1/17 (CD3+CD4+IFNγ+IL-17+), and Th1 cells (CD3+CD4+IFNγ+IL-17-) in collection media, consisting of 50% FBS and 50% phenol-free RPMI 1640 medium (11835030, ThermoFisher, Canada). Respective cell yields are provided in **Table S3**. Samples were stored on ice, added to respective DNA LoBind Tubes (EP022431021, Millipore Sigma, Canada), and centrifuged at 400RCF for 10mins at 4°C before resuspending pellets in 300uL of RNAprotect Cell Reagent (76526, Qiagen, Canada). Cells were stored at −80°C prior to RNA isolation using RNeasy Micro Kit (74004, Qiagen, Canada), as per manufacturer’s guidelines. RNA purity was assessed using the Nanodrop Spectrophotometer (ThermoFisher, Canada) prior to storage at −80°C. Th1, Th1/17, and Th17 populations from patients (n=10) and controls (n=12) were then sent for bulk RNA sequencing (RNAseq) at McGill Genome Centre (MGC, Canada) using Illumina NovaSeq X Plus 10B (Illumina, USA), generating approximately 30-51 million read pairs per sample, followed by subsequent bioinformatic analyses, as per standard MGC pipelines. Briefly, RNA samples were processed at MGC using the NEBNext® Ultra™ II Directional RNA Library Prep Kit (New England Biolabs, Canada) with poly(A) mRNA enrichment. Libraries were generated according to standard NEBNext protocols, including mRNA capture, fragmentation, first- and second-strand cDNA synthesis, end repair, adaptor ligation, and PCR amplification. Libraries underwent Ampure XP bead clean up and quality assessment (LibQC), including removal of primer dimer fragments, before pooling and sequencing using paired-end reads. Briefly, analyses included principal component analysis (PCA), unsupervised and supervised clustering, heatmap generation, differential expression analysis (DEG), volcano plots, fold-change distribution, and Gene Ontology (GO) enrichment analyses.

### Protein extraction from ectopic and eutopic tissue samples and multiplex cytokine array

Ectopic and eutopic tissues were immediately snap frozen upon collection and stored at −80°C prior to processing. Tissues were cut to a standardized weight of 37mg and placed in 1.4mm PowerBead tubes (13112-50, Qiagen, Canada) containing T-PER™ Tissue Protein Extraction Reagent (78510, ThermoFisher, Canada), as per manufacturer’s guidelines. Protein inhibitor cocktail (535140, Millipore Sigma, Canada) was added to each tube to reach a total volume of 400uL/tube. Samples were then mechanically digested using the Omni Bead Ruptor 24 (Omni International, USA), as per manufacturer’s guidelines, centrifuged at 10,000RCF for 5mins at 4°C, and protein extracts were aspirated. Protein concentrations were evaluated using a Pierce™ BCA Protein Assay Kit (23225, ThermoFisher, Canada), as per manufacturer’s instruction, and normalized to lowest concentration (1,202μg/mL). Protein extracts were then stored at −80°C prior to evaluation using a commercially available multiplex cytokine array (48-plex, EveTechnologies, Canada).

### Peritoneal fluid processing for multiplex cytokine array

PF was collected and placed on ice prior to centrifugation at 400RCF for 5mins at 4°C. Supernatant was aspirated, aliquoted, and stored at −80°C prior to multiplex cytokine array (48-plex, EveTechnologies, Canada). PF cells were cryopreserved (90% FBS and 10% DMSO) and stored at −80°C for later analysis. Prior to staining for flow cytometry (panel 3), PF cells were first thawed in a water bath at 37°C. Cell suspensions were added to warm RPMI media with 10% FBS and centrifuged at 400RCF for 10mins at 20°C prior to resuspension in warm TexMACS complete media and cell counting. PF cells were then seeded in 96-well plates at 1×10^6^ cells/well and re-stimulated prior to flow cytometry using 30ng/mL of PMA and 1μg/mL of ionomycin. After 1h of incubation, 2μL/mL of protein transport inhibitor (00-4980-93, ThermoFisher, Canada) was added for the last 3h of incubation (4h total) to allow for intracellular cytokine detection. Cells were then aspirated for flow cytometric analysis.

### Flow cytometric analyses

Flow cytometric analysis of human samples was conducted as detailed in our previous publication^5^, unless otherwise stated. Briefly, cells were first counted and stained with Human TruStain FcX™ (422302). All products were purchased from BioLegend, USA unless stated otherwise. PBMCs and PF cells were stained with extracellular Abs, as indicated in **Table S2**. To assess cell viability, PBMCs from panel 1 were stained with eBioscience™ Fixability Viability Dye eFluor™ 780 (65-0865-18, ThermoFisher, Canada), PBMCs from panel 2 were stained with Zombie Aqua™ Fixable Viability Kit (423101), and PF samples from panel 3 were stained with Zombie UV™ Fixable Viability Kit (423107). Following fixation and permeabilization, cells were stained with intracellular Abs, as indicated in **Table S2**. Cells were then washed. Data for panel 1 was acquired using the CytoFlex S (Beckman Coulter, USA), panel 2 was acquired using BD FACSAria™ III (BD Biosciences, USA), and panel 3 (PF cells) was acquired using BD FACSymphony™ (BD Biosciences, USA). Data was analyzed using FlowJo v10 software (Oregon, USA). UltraComp eBeads™ Compensation Beads (01-2222-42, ThermoFisher, Canada) were used to conduct compensation. FMO controls were used appropriately to determine gating strategy and an aliquot of half live and half heat-killed cells stained with viability dye was used to set a viability gate.

### Statistics

GraphPad Prism10®, USA was used for statistical analyses. When comparing two groups, in cases of normal (Gaussian) distribution, an unpaired Welch’s t test was performed. Whereas a Mann-Whitney test was conducted for unpaired, non-parametric data (two groups). For paired, non-parametric data, the Wilcoxon signed-rank test was performed (two groups). When comparing multiple groups, a two-way ANOVA was used followed by Šídák’s multiple comparisons test. Outliers were identified using the ROUT method (Q=1%) in GraphPad Prism and excluded prior to statistical analysis, as indicated in respective figure captions. Error bars indicate standard deviation (SD) and statistical significance was signified by a p-value < 0.05.

For RNAseq figures, genes with an adjusted p-value (FDR; False Discovery Rate) ≤ 0.05 were considered differentially expressed; no additional log2 fold change threshold was applied. All bioinformatic analyses were conducted by the McGill Genome Centre. Briefly, GenPipes was used for differential expression analysis as the main in-house framework of the C3G, used to perform major processing steps^25^. Adaptor sequences and low-quality score bases (Phred score <30) were trimmed from reads using Trimmomatic^26^. For context, Phred score (Q-score) is a quantitative measure of the base quality in DNA sequencing, whereby higher scores reflect greater confidence that the base was correctly identified. Resulting reads were aligned to the genome using STAR^27^. Samples with poor alignment metrics (i.e. <40% uniquely mapped reads) were excluded from downstream analysis to avoid artefactual differential expression. This filtering step reduces false positive DEGs that can occur when low-quality samples artificially inflate fold changes due to low total mapped reads. Read counts were obtained using HTSeq^28^. The R package DESeq2^29^ was used to identify differences in expression levels between groups using negative Binomial GLM fitting and Wald statistics: nbinomWaldTest. Additionally, “ashr”^30^ was used to shrink log2 fold changes in gene expression data to improve effect size estimation stability. For GO analysis, all genes which were differentially expressed were considered for input. Functional enrichment analyses (Gene Ontology) was performed using clusterProfiler^31^. P-values were obtained using a hypergeometric test with the total set of genes as background, and Benjamini-Hochberg FDR correction was applied for multiple testing.

### Study approval

The Queen’s University Health Sciences and Affiliated Teaching Hospitals Research Ethics Board approved all methodologies used in this study (OBGY-229-11 and ANAT-029-09). All participants provided written, informed consent prior to the collection of samples.

### Data availability

Main data supporting the findings of this study are available within this article and supplementary files. RNAseq data generated in this study has been deposited in the NCBI Gene Expression Omnibus (GEO) under accession number GSE313775.

## Results

### Key cytokines and chemokines involved in lesion establishment/maintenance are significantly dysregulated in EMS patient plasma, tissues, and peritoneal fluid

To assess systemic and local alterations in cytokine and chemokine levels, multiplex cytokine array was conducted using EMS patient and control plasma, EMS patient PF, and patient eutopic and ectopic tissues, analyzing levels of 48 key immunological analytes (**Figures 1 & 2**). IL-1β, IL-6, IL-8, and IL-18 analytes were significantly increased in patient plasma compared to controls (**Fig. 1A-D**). In contrast, G-CSF was significantly decreased in patient plasma as compared to healthy controls (**Fig. 1E**). Other analytes of interest due to their known roles in EMS pathophysiology and/or the IL-23/Th17/IL-17 axis, such as IL-17A, IL-23 (p40 subunit), and IFNγ, were not significantly altered in plasma (**Fig. 1F-H**). Notably, when results were stratified by disease stage, IL-6 levels were significantly increased in severe-stage (III-IV) relative to mild-stage (I-II) EMS (**Fig. 1I**).

**Figure 1:**
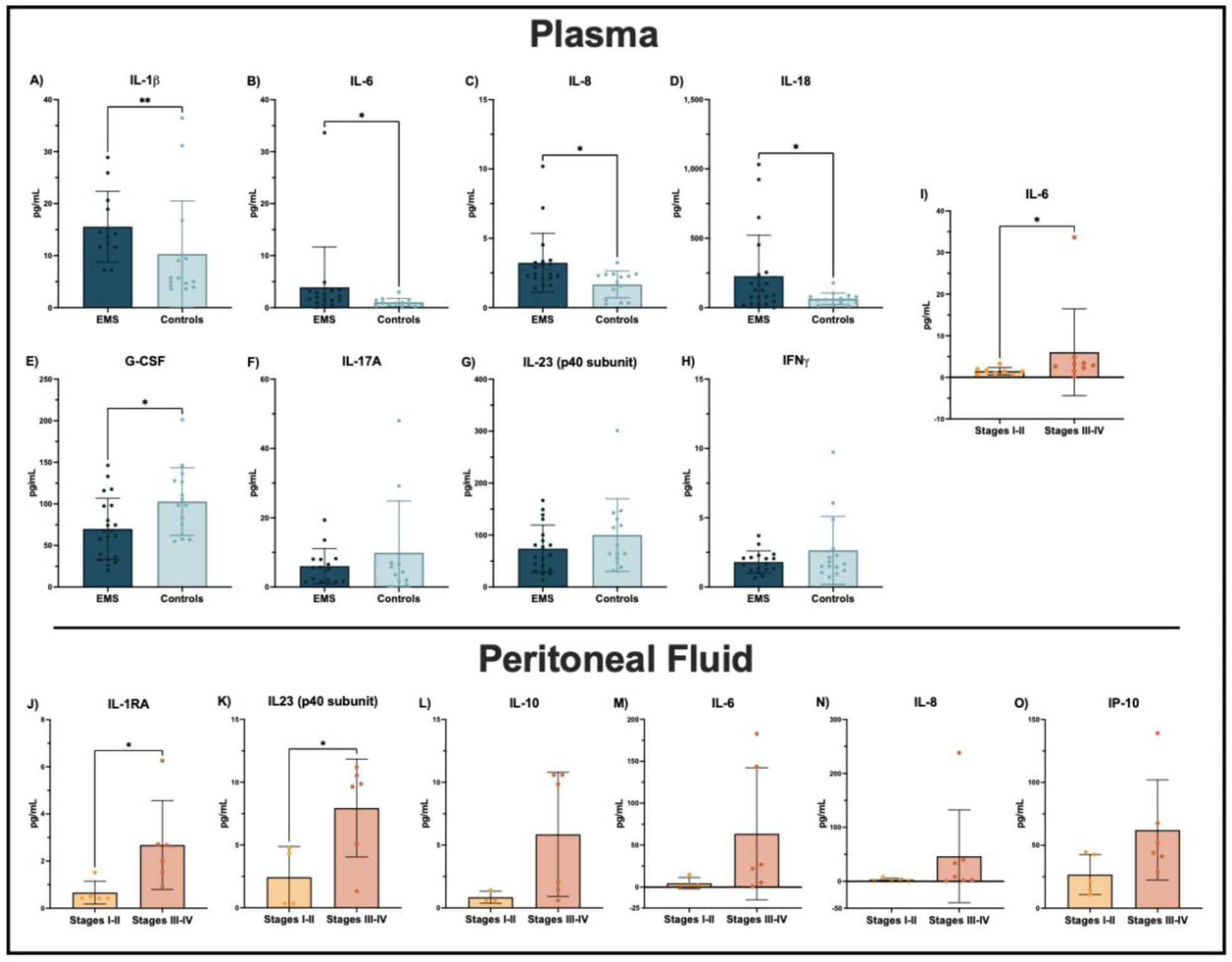
EMS patients have significantly altered levels of key immunological factors in plasma and PF compared to healthy controls, depicting systemic and local immune dysfunction in severe-staged disease. EMS patient (n=21) and control (n=15) plasma were collected and analyzed using a multiplex cytokine array. IL-1β, IL-6, IL-8, and IL-18 analytes were significantly increased in patient plasma (A-D), whereas G-CSF was significantly decreased in EMS patient plasma as compared to controls (E). Levels of other analytes of interest, such as IL-17A, IL-23 (p40 subunit), and IFNγ, were not significantly altered (F-H). When stratified by stage, plasma IL-6 levels were significantly increased in EMS patients with severe-staged (n=9; stages III-IV) relative to mild-staged (n=8, stages I-II) disease (I). Moreover, IL-1RA and IL-23 (p40 subunit) were significantly increased in the PF of patients with severe-staged (n=6; stages III-IV) relative to mild-staged (n=5, stages I-II) disease (J, K). Other key analytes of interest, such as IL-10, IL-6, IL-8, and IP-10, were not significantly altered between disease stages (L-O). Data is represented as mean ± SD. Statistical analyses were performed using the Mann-Whitney test for unpaired, non-parametric data. One outlier was identified in panel B using the ROUT method (Q=1%) and was retained in the analysis as biologically plausible. *p<0.05, **p<0.01.

When stratifying patient PF by disease stage, IL-1RA and IL-23 (p40 subunit) levels were significantly increased in severe-relative to mild-staged EMS (**Fig. 1J, K**). Levels of other analytes of interest, including IL-10, IL-6, IL-8, and IP-10, did not show statistically significantly differences (**Fig. 1L-O**). Additionally, results revealed that MIP-1α was significantly increased in patient eutopic tissues as compared to matched ectopic tissues (**Fig. 2A**). Other analytes of interest, such as VEGF, G-CSF, GROα, IL-6, IL-8, IL-17A, and IL-23 (p40 subunit), were not significantly altered in patient tissues (**Fig. 2B-H**). However, when results were stratified by EMS stage, eutopic tissues had significant stage-dependent differences in analyte levels. Namely, FLT-3L and G-CSF levels were significantly increased in the eutopic tissues of severe-staged patients (**Fig. 2I, J**), while RANTES was significantly increased in that of mild-staged patients (**Fig. 2K**).

**Figure 2:**
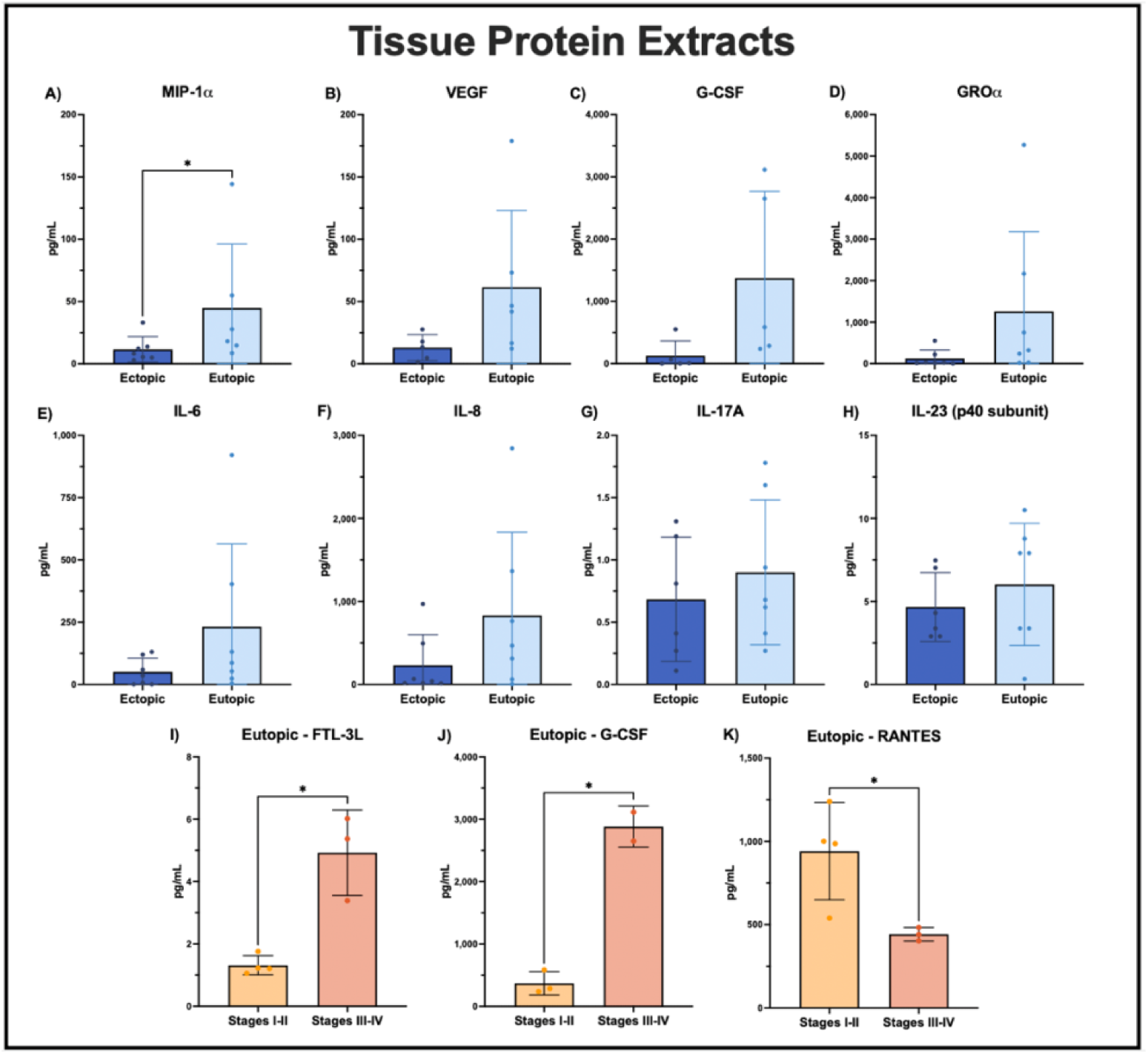
Protein extracts reveal significant alterations in the local EMS lesion microenvironment as compared to eutopic patient tissues. Briefly, matched patient eutopic and ectopic tissues (n=7) were collected for protein extraction. Cytokine/chemokine analyses revealed MIP-1α to be significantly increased in eutopic tissues of patients as compared to matched ectopic tissues, highlighting immune dysregulation in the eutopic endometrium (**A**). Other analytes of interest, including VEGF, G-CSF, GROα, IL-6, IL-8, IL-17A, and IL-23 (p40 subunit), were not significantly altered in patient tissues (**B-H**). When stratified by stage, FLT-3L and G-CSF levels within eutopic tissues were significantly increased in severe-staged (n=3; stages III-IV) relative to mild-staged (n=4; stages I-II) disease (**I, J**). Whereas RANTES levels were increased in eutopic tissues of those with mild-stage disease (**K**). Data is represented as mean ± SD. Statistical analyses were performed using the Wilcoxon signed-rank test for paired, non-parametric data (A-H) or an unpaired Welch’s t test for independent samples (I-K). *p<0.05.

### EMS patients exhibit significantly altered frequencies of key T cell subsets in peritoneal fluid

To gain a comprehensive overview of Th17 and Treg cell subsets within the local lesion microenvironment, we then conducted high-parameter flow cytometry on patient PF (**Figure 3A**). Results showed that EMS patients have significantly increased pTh17 cells as compared to both Tregs and npTh17 cells (**Fig. 3B, C**). Moreover, when looking broader at total Th17 cells, this population was still significantly increased as compared to Tregs (**Fig. 3D**). Though, frequency of total PF Th17 cells did not significantly differ between mild- and severe-stage EMS patients (data not shown). Additionally, while no stage-specific differences were found in IFNγ expression on pTh17 cells (**Fig. 3E**), it was apparent that EMS patients have significantly decreased frequency of CCR6+ Th17 cells, with almost all Th17 cells being CCR6-(**Fig. 3F**).

**Figure 3:**
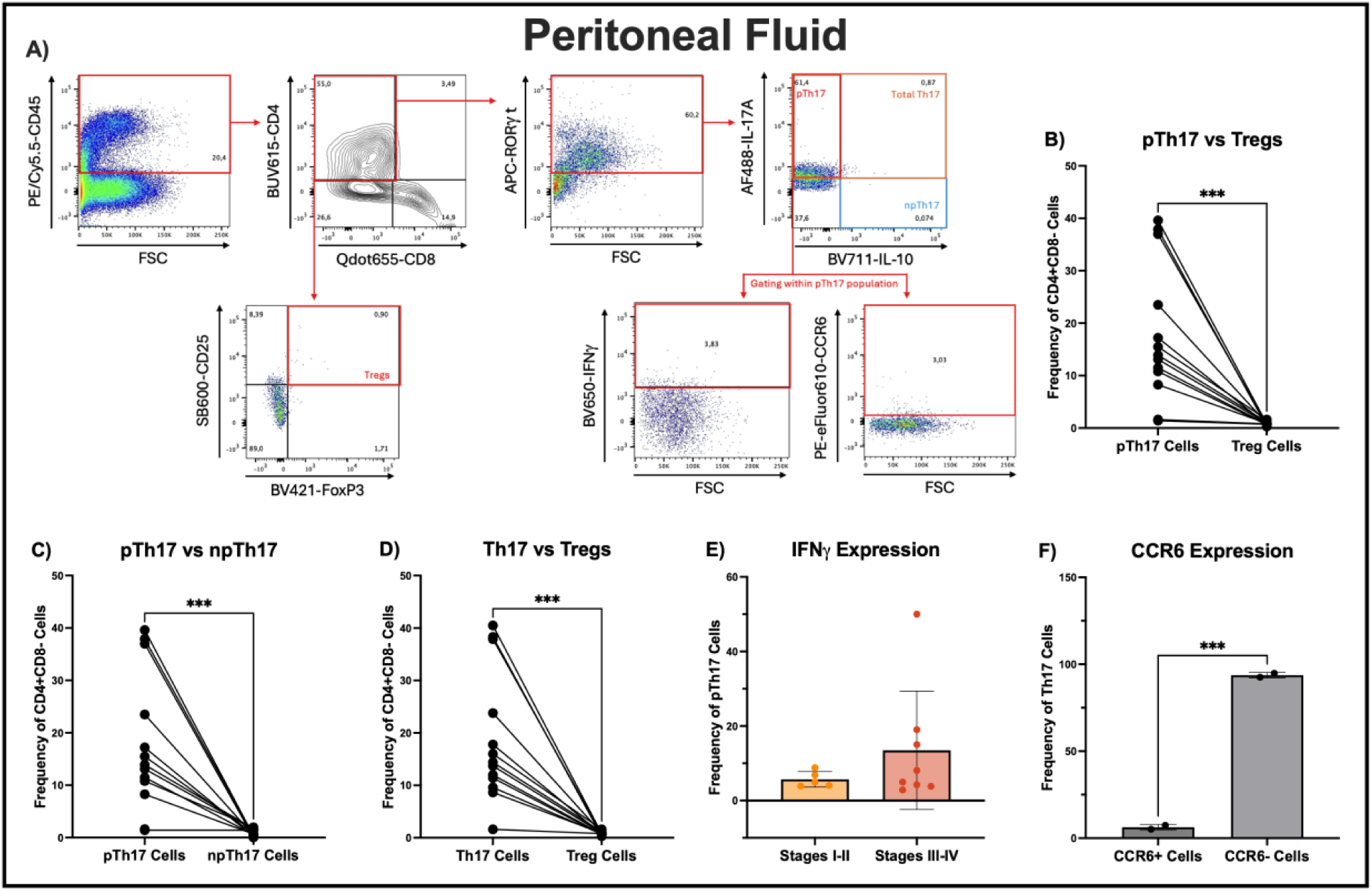
EMS patients have significantly different frequencies of key T cell subsets within PF. EMS patient PF cells were analyzed using flow cytometry, as per the shown gating strategy (**A**). Shown population frequencies (%) represent cell proportions within each respective parent gate/population. Results show that patients (n=13) have significantly increased pTh17 cells (CD45+CD8-CD4+RORγt+IL-17A+IL-10-) as compared to both Treg (CD45+CD8-CD4+CD25+FoxP3+) and npTh17 (CD45+CD8-CD4+RORγt+IL-17A-IL-10+) cell subsets. (**B, C**). Moreover, total Th17 cells (CD45+CD8-CD4+RORγt+IL-17A+) were significantly increased within the CD45+CD4+CD8-population as compared to Tregs (**D**). Frequencies of these T cell subsets did not significantly differ when stratified by disease stage. Moreover, there were no significant, stage-dependent differences in IFNγ expression within the pTh17 population (**E**). When assessing CCR6 expression, patients were found to have a significantly increased CCR6-population of Th17 cells (**F**). Data is represented as mean ± SD. Statistical analyses were performed using either the Wilcoxon signed-rank test for paired, non-parametric data (B-D, F) or a Mann-Whitney test for unpaired, non-parametric data (E). One statistical outlier was identified and excluded from the analysis in panel C using the ROUT method (Q=1%). ***p<0.001.

### An IL-23-containing Th17 cocktail and activation promotes a significant increase of Th17 cells from patient PBMCs

We then sought to determine the effect of *in vitro* stimulation of patient/control PBMCs with an IL-23-containing Th17 cocktail, assessing the impact on Th17 cell frequency after 4 days in culture using flow cytometry (**Figure S1A**). When in combination with CD3/CD28 activation, this Th17 cocktail promoted the most significant increase in Th17 cell frequency from patient PBMCs, as compared to those treated with Th17 cocktail and no activation as well as the media control (**Fig. S1B**). These significant alterations in Th17 frequency were not observed in control PBMCs.

### EMS patients have significant changes in circulating Th17 and Th1 cells proportions, along with distinct transcriptional profiles of circulating Th1, Th1/17, and Th17 cells

To assess systemic changes in key T cell subsets (Th1, Th1/17, and Th17), we first sorted these populations from patient and control PBMCs using the FACSAria III (**Figure 4A**). Sorting data revealed no significant differences in the total number of circulating Th17 and Th1/17 cells (**Fig. 4B, C**). However, circulating Th1 cells were significantly decreased in patients as compared to controls (**Fig. 4D**). When results were stratified by EMS stage, patients had significantly increased Th17 cells in mild-staged disease (stages I-II) compared to those in severe-staged disease (stages III-IV; **Fig. 4E**). There were no statistically significant stage-dependent differences in Th1/17 and Th1 cell subsets (**Fig. 4F, G**).

**Figure 4:**
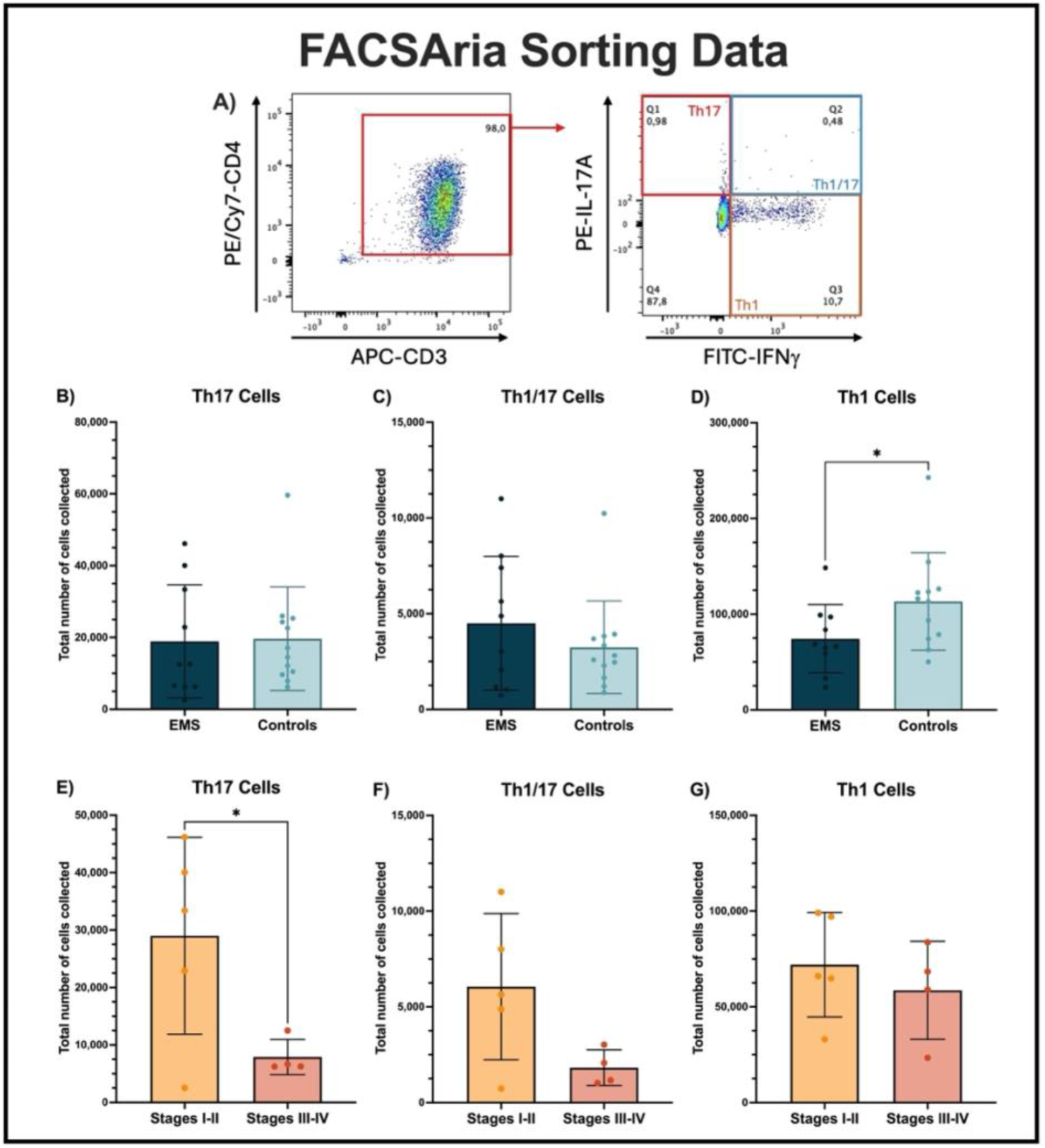
EMS patients have significantly decreased Th1 cells in circulation and significantly increased circulating Th17 cells in mild-staged disease. Briefly, patient (n=10) and control (n=12) PBMCs were sorted using FACSAria III (**A**). While there were no significant differences in total number of circulating Th17 (CD3+CD4+IFNγ-IL-17A+) or Th1/17 (CD3+CD4+IFNγ+IL-17A+) cell subsets (**B, C**), circulating Th1 cells (CD3+CD4+IFNγ+IL-17A-) were significantly decreased in patients compared to controls (**D**). When stratified by stage, EMS patients had significantly increased Th17 cells in mild-staged (n=5; stages I-II) relative to severe-staged (n=4; stages III-IV) disease (**E**), whereas there were no significant differences in Th1/17 and Th1 cell subsets when staged (**F, G**). Data is represented as mean ± SD. Statistical analyses were performed using an unpaired Welch’s t test. *p<0.05.

Bulk RNAseq was then performed to comprehensively characterize transcriptional profiles of Th1, Th1/17, and Th17 cell subsets isolated from patients and controls (**Figures 5-7**). Results depict that Th17 cells have a unique transcriptional profile in EMS patients as compared to controls, with 2,220 DEGs (2074 upregulated and 146 downregulated) identified (**Fig. 5A**). The top 5 upregulated DEGs were *MUC3A* (fold change: 9.3)*, HYDIN2* (9.2)*, CACNA1S* (9.0)*, TF* (8.7), and *C2CD5* (8.6) and top 5 downregulated DEGs were *PLS3* (−5.3)*, C1QC* (−5.3)*, H1-2* (−4.8)*, HLA-DMA* (−4.0), and *SERPINE1* (−3.6; **Table S4**). Broadly, this profile points to EMS patients having Th17 cells with heightened potential for inflammation and altered functional properties, which may contribute to associated chronic inflammation and tissue remodeling in EMS. Notably, Th17 cells also had significantly downregulated *IL-17F* (−2.8) and *IL-17A* (−2.8) in patients as compared to controls, possibly indicating a dysfunctional or exhausted phenotype due to chronic antigen exposure. As depicted by a supervised heatmap, there is a clear distinction in transcriptional profile of Th17 cells from EMS patients and controls, with EMS-derived Th17 cells showing a broad upregulation of genes, highlighting a unique transcriptional signature (**Fig. 5B**). GO and KEGG enrichment analyses depict DEGs to be strongly associated with ion channel activity and transport, membrane-associated processes, neuronal and sensory system processes, and cardiac and calcium signaling pathways (**Fig. 5C**).

**Figure 5:**
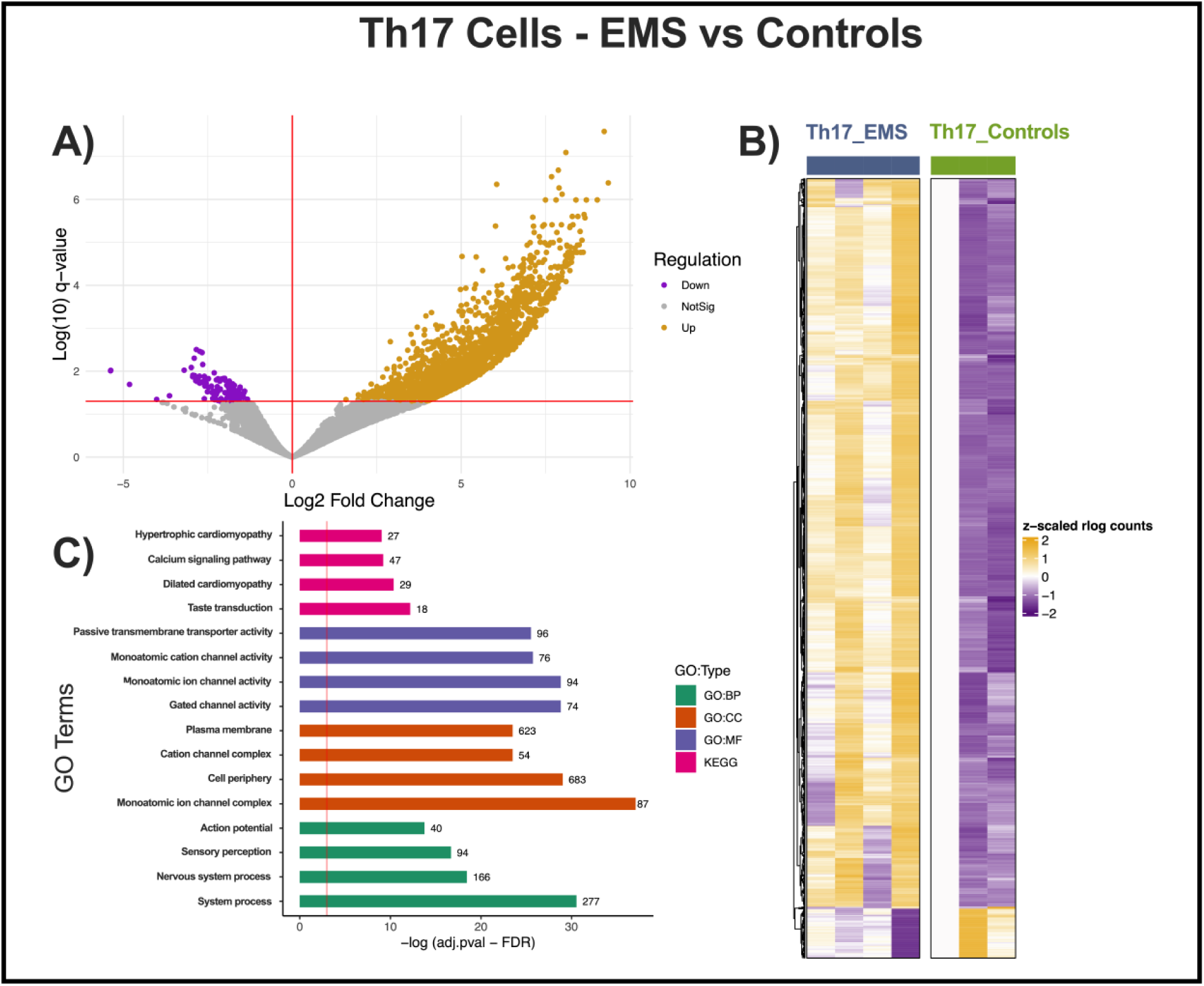
Th17 cells have a unique transcriptional profile in EMS patients compared to healthy controls. A volcano plot depicts 2,220 DEGs (2074 upregulated and 146 downregulated) in Th17 cells of EMS patients compared to controls (**A**). A supervised heatmap depicts Th17 gene expression patterns in each cohort (**B**). A GO plot depicts enrichment of specific GO terms within significant DEGs (**C**). Green (GO:BP) = Biological Process, Orange (GO:CC) = Cellular Component, Purple (GO:MF) = Molecular Function, Pink (KEGG) = KEGG pathways.

When comparing Th1 cells in patients and controls, 42 DEGs (39 upregulated and 3 downregulated) were discovered (**Fig. 6A**). The top 5 upregulated DEGs were *PARD6G* (fold change: 3.0), NA (novel transcript, 3.0), *ZNF133* (2.9)*, ADGRG5* (2.9), and *WWTR1* (2.5) and top 3 downregulated DEGs were NA (novel transcript, −3.2), *ZNG1F* (−1.8), and *ROCK1P1* (−1.3, **Table S5**). Generally, upregulated genes suggest that patients may have altered polarity, signaling, and survival of Th1 cells, as well as a slight downregulation of regulatory/cytoskeletal-associated genes. Interestingly, *MUC1* (1.4) was significantly upregulated in EMS patient Th1 cells compared to controls, suggesting an aberrant, disease-specific transcriptional program. A supervised heatmap also depicts Th1 gene expression patterns in EMS patients compared to controls (**Fig. 6B**). Furthermore, GO analysis revealed DEGs to be most strongly associated with the Molecular Function category, with “molecular transducer activity” and “signaling receptor activity” being the most significantly enriched terms (**Fig. 6C**). Ultimately, compared to Th17 data, Th1 cells depict a more subtle transcriptional shift in EMS patients, suggesting that Th17 dysregulation may contribute more significantly. Though, further work is required to determine if Th17 dysregulation is a result of EMS or a driving force in disease pathophysiology.

**Figure 6:**
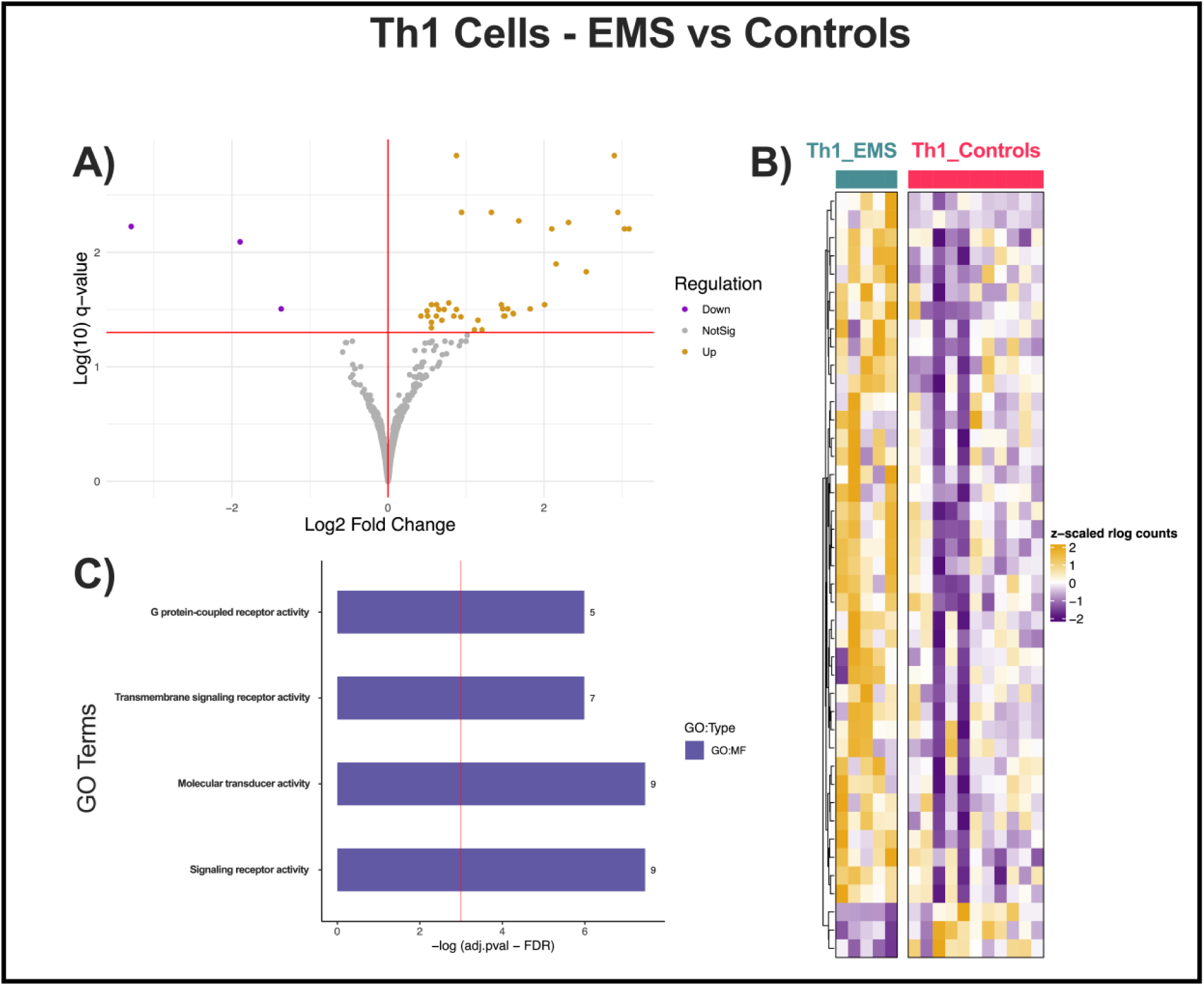
Th1 cells have a unique transcriptional profile in EMS patients compared to healthy controls. A volcano plot depicts 42 DEGs (39 upregulated and 3 downregulated) in Th1 cells of EMS patients compared to controls (**A**). A supervised heatmap depicts Th1 gene expression patterns in each cohort (**B**). A GO plot depicts enrichment of specific GO terms within significant DEGs (**C**). Purple (GO:MF) = Molecular Function.

When comparing T cell subsets within the EMS patient cohort, significant differences were again found in transcriptional profiles (**Figures 7, S2, & S3**). Upon comparing Th17 and Th1 cells in EMS patients, we identified 1,573 DEGs (1,512 upregulated and 61 downregulated, **Fig. 7A**). The top 5 upregulated DEGs were *IL17A* (9.4)*, LDB3* (7.1)*, REELD1* (7.0)*, SHISA6* (6.9), and *IL17F* (6.7) and top 5 downregulated DEGs were *IFNG* (−7.3), *NSF* (−5.4)*, XCL1* (−4.7)*, RNPEP* (−4.7), and *CCL4* (−4.7, **Table S6**). This aligns with literature depicting that Th17 and Th1 cells are major producers of IL-17A/F and IFNγ, respectively. A supervised heatmap depicts a clear distinction in transcriptional profiles of Th17 and Th1 cells compared in patients, with Th17 cells generally showing a broader upregulation of genes (**Fig. 7B**). Moreover, a distribution plot of gene expression changes comparing Th17 and Th1 cells depicts more pronounced transcriptional differences in EMS compared to controls, with an increased number of genes upregulated in Th17 cells (**Fig. 7C**). This may suggest a stronger Th17-driven signature in EMS as compared to healthy controls. Significant differences were also seen when comparing Th1/17 and Th17 cells (**Fig. S2 & Table S7**), as well as Th1/17 and Th1 cells (**Fig. S3 & Table S8**) within the patient cohort, as again depicted by volcano plots, heatmaps, and GO/KEGG analyses.

**Figure 7:**
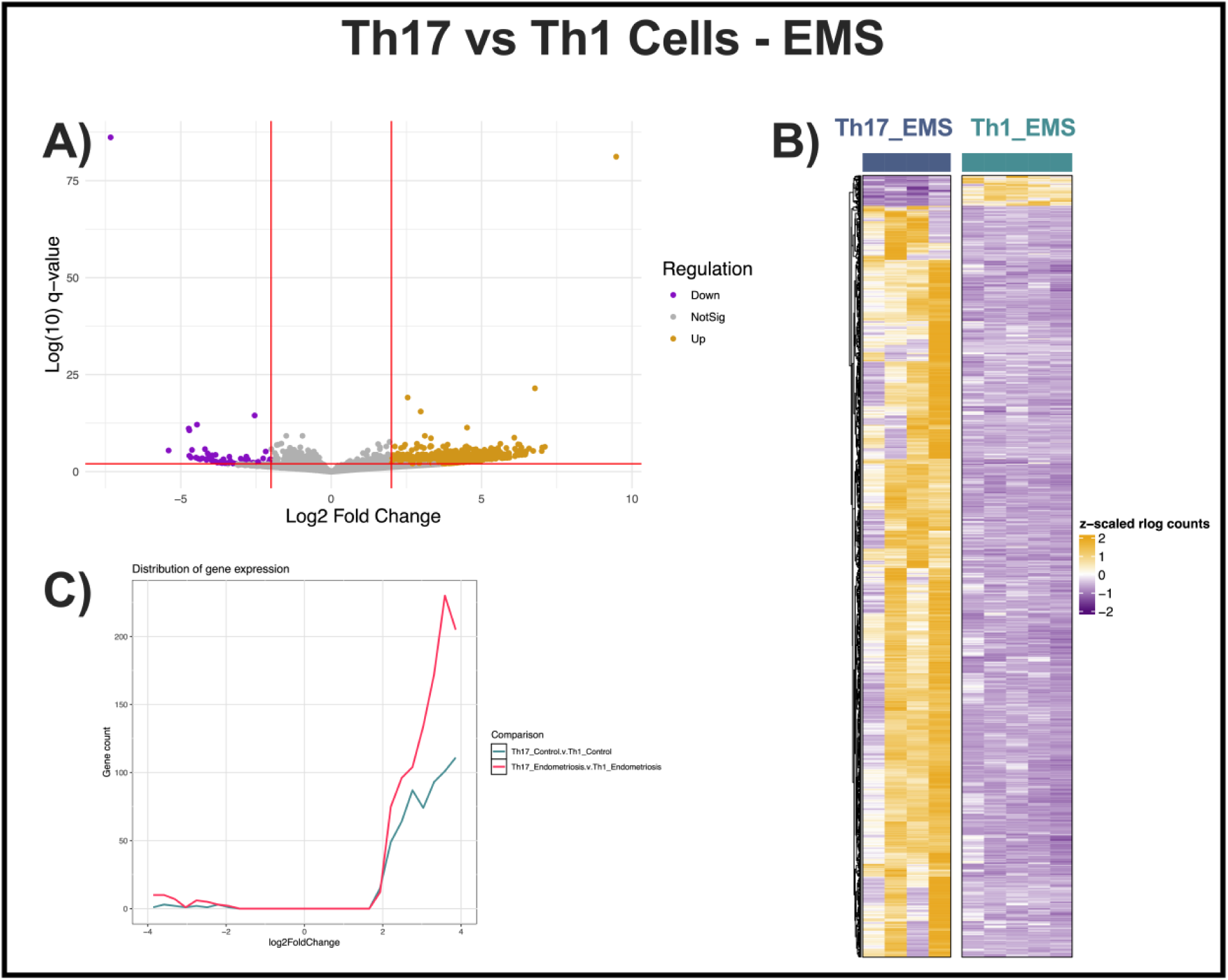
Within EMS patients, Th17 and Th1 cell subsets have significant differences in transcriptional profiles. A volcano plot depicts 1,573 DEGs (1,512 upregulated and 61 downregulated) in Th17 cells compared to Th1 cells within EMS patients (**A**). A supervised heatmap depicts Th17 vs Th1 gene expression patterns in patients (**B**). A plot of fold change density per comparison depicts a larger distribution of gene expression when comparing Th17 vs Th1 cell subsets in patients (red) compared to controls (blue; **C**).

## Discussion

The EMS lesion microenvironment is a complex, heterogeneous system wherein there is a dynamic interplay between immune and non-immune (epithelial, stromal, endothelial) cell subsets. Evidence in literature and our previous work suggests that sterile inflammation in EMS is largely driven by a dysfunctional immune system. Indeed, altered T cell subsets, including Tregs and Th17 cells, are reported to be dysfunctional in EMS, promoting chronic inflammation and dampening immune clearance of lesions^5,7,16^. While exact mechanisms are still unknown, we and others have shown that the IL-23/Th17/IL-17 axis is dysregulated in EMS and likely contributes to the disease pathophysiology^4–9^. However, it is still not understood whether this immune dysfunction is driving EMS, or if this is a consequence of disease. To gain a better understanding of the complex interplay between Th subsets in EMS pathology, we examined systemic and localized changes specifically to gain a comprehensive understanding of how these Th subsets are reprogrammed in EMS and contribute to disease.

Using patient plasma, PF, and protein extracts from matched eutopic and ectopic tissues, we observed significant local and systemic alterations in key cytokines/chemokines known to drive EMS lesion establishment and maintenance. Indeed, EMS patients exhibited markers of systemic inflammation, as characterized by significantly increased circulating levels of IL-1β, IL-6, IL-8, and IL-18 in EMS plasma compared to controls. In contrast, G-CSF levels were significantly decreased in patient plasma. When stratified by disease stage, IL-6 was significantly increased in severe-stage (stages III-IV) relative to mild-stage (stages I-II) EMS, depicting that IL-6 is not only increased in patients compared to controls, but also associated with EMS disease severity. This is in accordance with previous literature^32,33^.

In the local EMS lesion microenvironment, we report significantly decreased MIP-1α within patient ectopic tissues compared to eutopic tissues. As MIP-1α is known to promote activation/recruitment of immune cells and tissue repair^34^, this suggests a dysregulated immune microenvironment in ectopic tissues, likely allowing for immune evasion and lesion persistence. When stratified by disease stage, FLT-3L and G-CSF levels were significantly increased in the eutopic tissues of severe-stage relative to mild-stage patients, further suggesting that eutopic endometrium from EMS patients harbours an inflammatory microenvironment. This is in line with previous reports suggesting specific alterations within EMS patient eutopic endometrium associated with EMS pathophysiology^35^. In contrast, RANTES was significantly increased in eutopic tissues of mild-stage relative to severe-stage EMS. Taken together, multiplex cytokine array results reveal stage-specific alterations of the eutopic endometrial immune microenvironment in EMS. As MIP-1α, FLT-3L, and RANTES are known to be produced by T cells^36,37^, and evidence suggests T cell dysfunction in EMS, altered Th subsets may be responsible for driving these local and systemic modifications to immune mediators in EMS^38^.

Within patient PF, results depict localized increases of IL-1RA and IL-23 (p40 subunit) in severe-stage EMS relative to mild-stage patients, highlighting that these immune mediators may be associated with disease severity. Though, it should be noted that this p40 subunit of IL-23 is also shared with IL-12, and as such, further analyses are necessary to determine if this variance is specific to IL-23. High-parameter flow cytometry revealed significantly increased pTh17 cell frequency within CD4+CD8-cells in EMS patients as compared to both Treg and npTh17 populations. Frequency of total Th17 cells was also significantly increased in EMS patients compared to Tregs. Within this Th17 population, patients had significantly more CCR6-cells. As literature suggests that CCR6^low^/CCR6^neg^ Th17 cells tend to represent a non-classical or more plastic Th17 subset^39^, this may provide insight into how Th17 cells functionally differ in EMS. Moreover, as we report increased levels of IL-23 in severe-stage patient PF (and previously in patient plasma^5^), which is known to largely drive pTh17 cell fate^5,40^, as well as increased frequency of pTh17 cells in EMS patients, this may indicate a pTh17 dominant microenvironment. Though, as IL-1RA is known to suppress Th17 differentiation and promote Treg cell fate^41^, this may be increased in severe-stage patients in attempt to regulate Th17/Treg ratio and promote immune homeostasis.

Further systemic alterations were identified during sorting of patient PBMCs into Th17, Th1/17, and Th1 populations. Specifically, EMS patients had significantly decreased circulating Th1 cells as compared to healthy controls. When stratified by disease stage, circulating Th17 cells were significantly increased in mild-stage relative to severe-stage EMS. While Th17 cells have been associated with disease severity within patient PF^7^ and shown to be increased in patient peripheral blood^14^, stage-specific differences in circulating Th17 populations remain less well characterized. Our findings suggest that circulating and local Th17 populations may not follow identical patterns across disease stages, highlighting the complexity of systemic and lesion-associated immune responses in EMS.

Finally, using bulk RNAseq we were able to comprehensively characterize transcriptional profiles of circulating Th17, Th1/17, and Th1 cells in EMS patients. Overall, EMS-derived Th17 cells had extensive transcriptional reprogramming, with 2,220 DEGs as compared to more modest changes in Th1 cells (42 DEGs). These results suggest transcriptional alterations in Th17 cells in EMS and support evidence suggesting that Th17 cell dysfunction and/or plasticity likely play a key role in EMS pathology. In conjunction with findings of increased CCR6-Th17 cells in patients, downregulation of IL-17A/F in patient Th17 cells may indicate an aberrant Th17 cell phenotype in EMS. While we have not conducted functional studies to document plasticity, our findings suggest that repeated antigen exposure and immune system overload associated with delayed clearance of debris in EMS likely contribute to an exhausted Th17 cell phenotype or reveal an immunoregulatory effort to reduce overt IL-17 responses. Moreover, GO and KEGG analyses of sequencing data from Th17 cells reveal that, as compared to healthy controls, EMS-derived Th17 cells have enriched ion transport, membrane, and calcium signaling pathways, which are recognized to play a key role in neuro-immune crosstalk and pain processes in EMS. Furthermore, *MUC1* was significantly upregulated in EMS-derived Th17 cells as compared to healthy controls. This is notable as *MUC1* is commonly overexpressed in cancer, where it can inhibit T cell proliferation and function, creating an immunosuppressive environment^42^. In contrast, Th1 cells exhibited more subtle transcriptional changes, mainly enriched for signaling-related molecular functions. Broadly, these functional variations suggest altered receptor-mediated signaling in EMS-derived Th1 cells, possibly reflecting changes in their ability to respond to inflammatory/tissue-derived signals within the EMS lesion microenvironment.

Patient characteristics revealed significant differences between EMS patients and controls in hormone use at time of sample collection, as well as over the counter (OTC) medication use. Given the known immunomodulatory effects of hormonal therapies and certain OTC medications, these variables may have influenced the immune profiles observed in this study. However, these differences also reflect the clinical reality of EMS, where hormonal treatments and analgesic use are common components of disease management. In contrast, no significant differences were observed between groups with respect to comorbidity prevalence, prescription medication use, menstrual status, or days since the first day of the last menstrual period. These findings suggest that sample acquisition did not occur at systemically different points in the menstrual cycle between groups.

Due to the invasive nature of sample collection, we were unable to obtain tissue and PF samples from healthy control volunteers for multiplex cytokine analysis. While comparisons of eutopic and ectopic tissues can help us gain knowledge on the EMS lesion microenvironment and the eutopic endometrium, literature shows that eutopic tissues from patients exhibit signs of immune dysfunction^5,43^, and as such, this tissue is not a true control. Moreover, while we identified significant differences in Th subset frequencies in EMS patients, due to this limitation in obtaining control PF, it is unclear how these frequencies differ to that of healthy controls. Patient samples used in this study were staged using rASRM staging criteria to enable comparison of immune dysfunction between mild-stage (I-II) and severe-stage (III-IV) EMS. However, as aforementioned, this staging criteria has been criticized due to limited correlation with patient symptoms, such as pain and infertility. Thus, while results stratified by stage may help us gain insight on immune alterations correlated with EMS disease severity, this limitation should be acknowledged.

In conclusion, we report significant local and systemic immune dysfunction in EMS as well as significant alterations in function and frequency of key Th cell subsets, potentially driving immune dysfunction in EMS. We highlight how these immune mediators and Th cell subsets may be influencing chronic inflammation in EMS and lesion establishment/maintenance through various mechanisms. To our knowledge, for the first time we report unique transcriptional profiles of Th1, Th1/17, and Th17 cells in EMS patients, providing insights for future studies as well as possible therapeutic targets. Ultimately, through this study, as well as our previous work^5,6,44^, we have identified two possible therapeutic targets for EMS patients, IL-17 and IL-23.

## Author contributions

DS conceived and conducted experiments, analyzed data, and wrote the manuscript. KZ and JH conducted experiments. OB contributed human patient samples. JP and KK aided with study coordination and sample transport. CT contributed reagents, conceived experiments, and provided financial support. All authors read and edited the manuscript.

## Acknowledgements

We would like to thank Zhilin Chen and Jefferey Mewburn from the Dept. of Biomedical and Molecular Sciences (DBMS, Queen’s University) Core Research Services for aid with flow cytometry/FACS. We would also like to thank François Lefebvre, Gerado Zapata, and Dr. Tony Kwan from the McGill Genome Centre (MGC, McGill University) for RNAseq sample processing and bioinformatic analysis.

## Supplemental information index

**Document S1.** Figures S1-S3, Tables S1, S2, and S3.

**Table S4:** Fold change values for upregulated (n=2,074) and downregulated (n=146) DEGs from RNAseq analysis of Th17 cells from EMS patients compared to controls, related to Figure 5.

**Table S5:** Fold change values for upregulated (n=39) and downregulated (n=3) DEGs from RNAseq analysis of Th1 cells from EMS patients compared to controls, related to Figure 6.

**Table S6:** Fold change values for upregulated (n=1,512) and downregulated (n=61) DEGs from RNAseq analysis of Th17 and Th1 cells within the EMS cohort, related to Figure 7.

**Table S7:** Fold change values for upregulated (n=61) and downregulated (n=229) DEGs from RNAseq analysis of Th1/17 and Th17 cells within the EMS cohort, related to Figure S2.

**Table S8:** Fold change values for upregulated (n=370) and downregulated (n=5) DEGs from RNAseq analysis of Th1/17 and Th1 cells within the EMS cohort, related to Figure S3.

