## Supplementary material for "An investigation of the IL-23/Th17 axis and transcriptomic profiles of T helper subsets in endometriosis": Document S1

**SUPPLEMENTAL FIGURES**

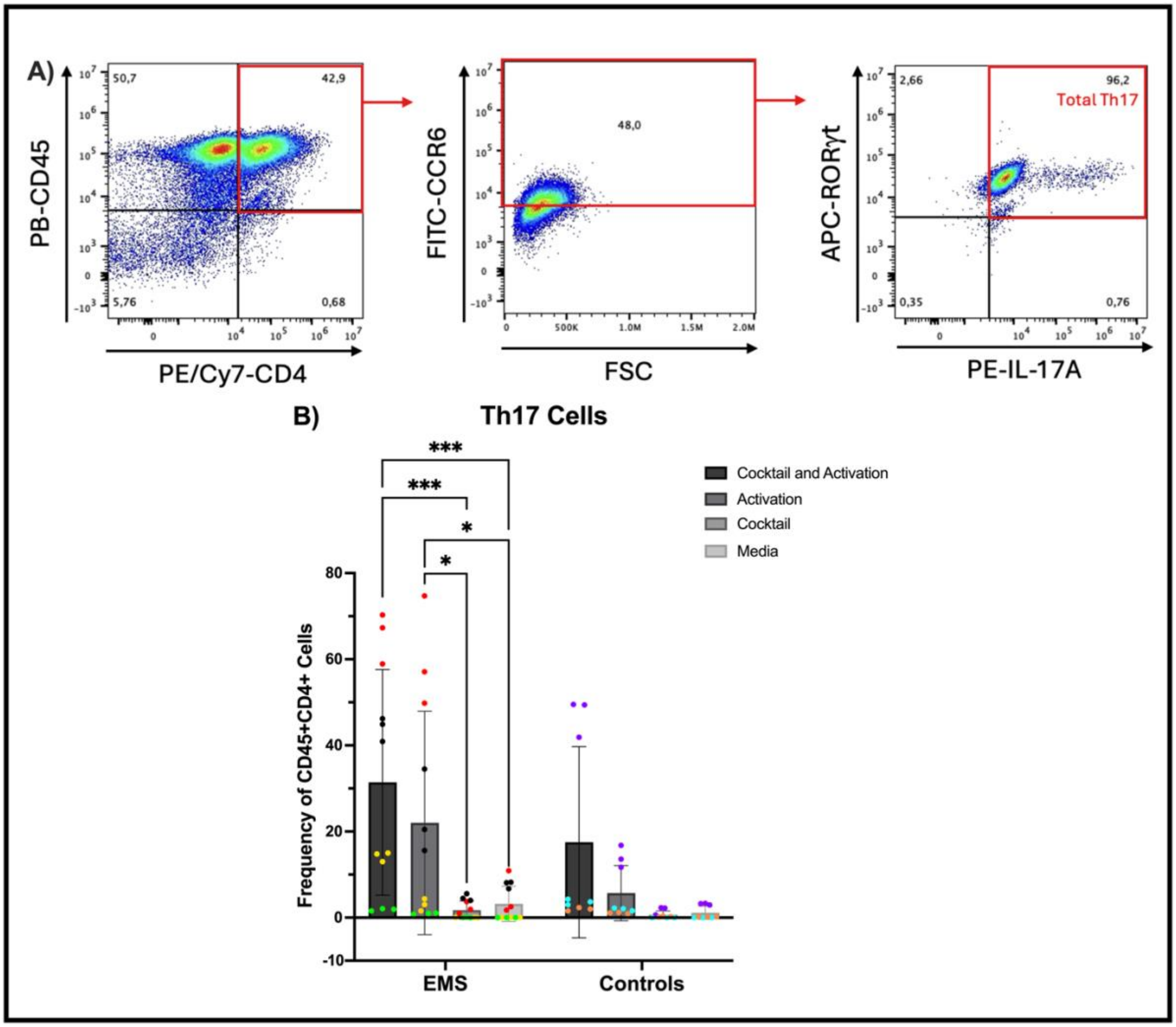

Figure S1: ***In vitro* stimulation of PBMCs with Th17 cocktail and activation promotes significantly increased expansion of Th17 cells in EMS patients**, as individually distinguished by coloured data points. PBMCs were first isolated from whole blood of EMS patients (n=4) and controls (n=3) and treated *in vitro* in triplicates with either i) Th17 cocktail and CD3/CD28 activation, ii) activation alone, iii) cocktail alone, iv) or media. Cells were incubated for 4 days prior to re-stimulation with PMA and ionomycin before flow cytometric analysis, in which, Th17 cells were classified as CD45+CD4+CCR6+RORγt+IL-17A+ cells (**A**). Stimulation with the Th17 cocktail and activation resulted in the greatest increase in Th17 cell frequency from patient PBMCs (**B**). Data is represented as mean ± SD. Statistical analyses were performed using two-way ANOVA followed by Šídák's multiple comparisons test. *p<0.05, ***p<0.001.

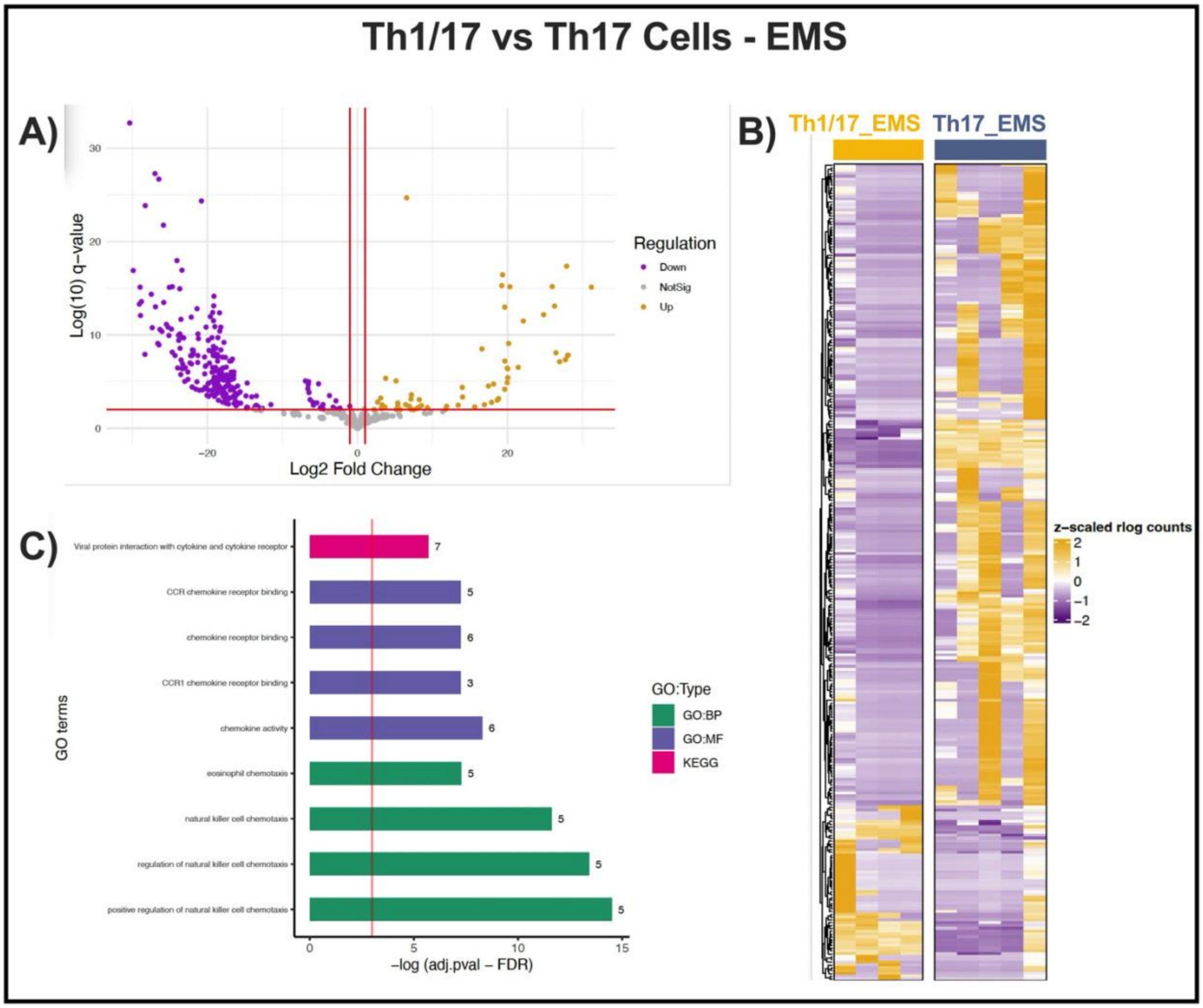

**<u>Figure S2:</u> Within EMS patients, Th1/17 and Th17 cell subsets have significant differences in transcriptional profiles.** A volcano plot depicts 290 DEGs (61 upregulated and 229 downregulated) in Th1/17 cells compared to Th17 cells within EMS patients (**A**). A supervised heatmap depicts Th1/17 vs Th17 gene expression patterns in patients (**B**). A GO plot depicts enrichment of specific GO terms within significant DEGs (**C**). Green (GO:BP) = Biological Process, Purple (GO:MF) = Molecular Function, Pink (KEGG) = KEGG pathways.

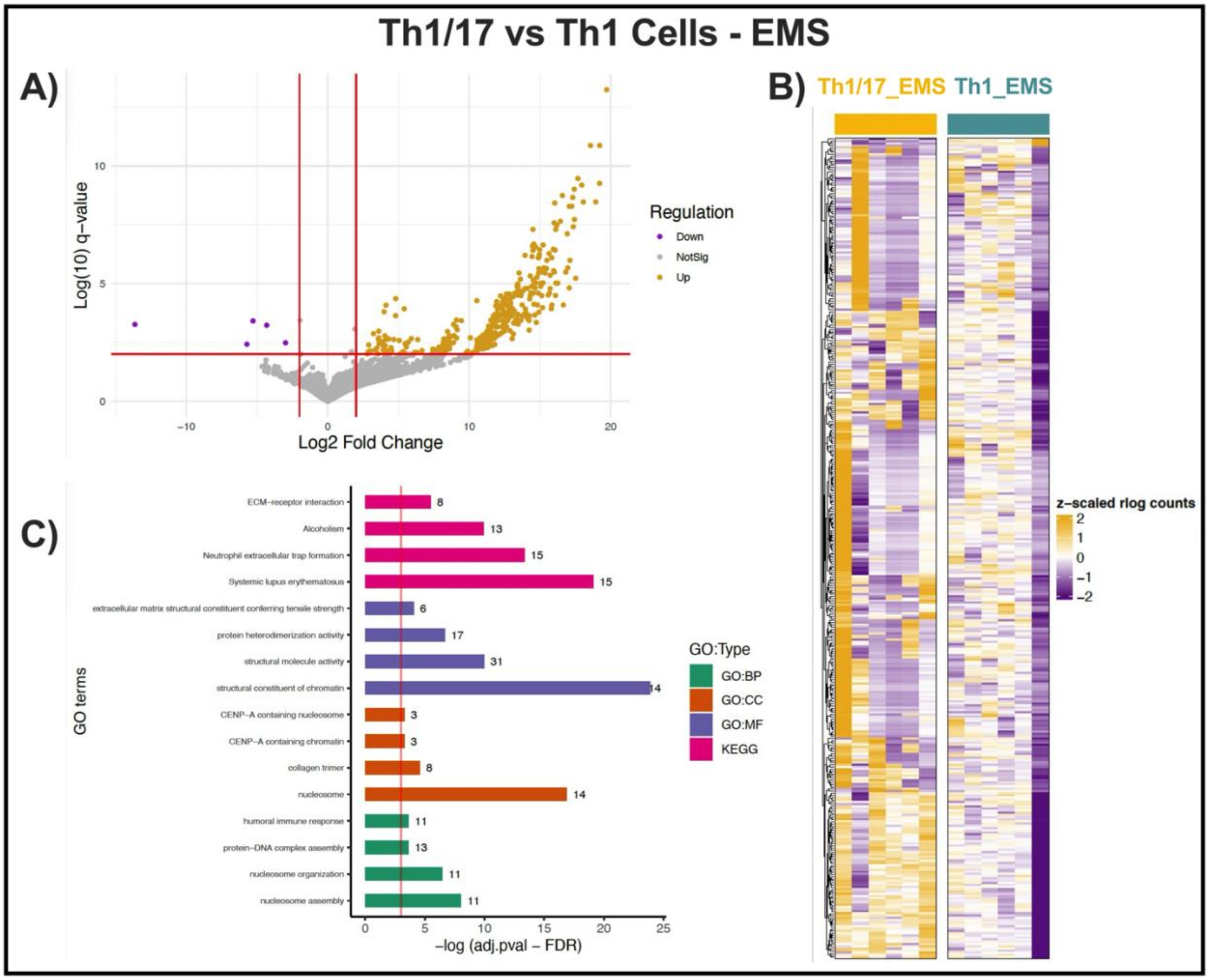

<u>Figure S3:</u> **Within EMS patients, Th1/17 and Th1 cell subsets have significant differences in transcriptional profiles.** A volcano plot depicts 375 DEGs (370 upregulated and 5 downregulated) in Th1/17 cells compared to Th1 cells within EMS patients (**A**). A supervised heatmap depicts Th1/17 vs Th1 gene expression patterns in patients (**B**). A GO plot depicts enrichment of specific GO terms within significant DEGs (**C**). Green (GO:BP) = Biological Process, Orange (GO:CC) = Cellular Component, Purple (GO:MF) = Molecular Function, Pink (KEGG) = KEGG pathways.

### TABLES

**<u>Table S1:</u> EMS patient and control study parameters, as indicated by self-questionnaire.** Statistical analyses were performed using a Fisher's exact test for all categorical variables, other than age, whereby a Welch's t test was used.

| Parameters | EMS (n=22): | Controls (n=19): | P-value: |
|---|---|---|---|
| Age ± SD | 38.1 ± 5.2 | 38.6 ± 2.9 | 0.8162 |
| *Post-secondary education, n (%)* | | | |
| Yes | 20 (90.9) | 19 (100.0) | 0.4902 |
| Other/None | 2 (9.1) | 0 (0.0) | |
| *Race, n (%)* | | | |
| Caucasian | 19 (86.4) | 16 (84.2) | 0.8145 |
| East Asian | 0 (0.0) | 1 (5.3) | |
| Other | 3 (13.6) | 2 (10.5) | |
| *Ever pregnant, n (%)* | | | |
| Yes | 10 (45.5) | 11 (57.9) | 0.8516 |
| No | 9 (40.9) | 6 (31.6) | |
| Unknown | 1 (4.5) | 0 (0.0) | |
| No response | 2 (9.1) | 2 (10.5) | |
| *Stage of EM, n (%)* | | | |
| I | 7 (31.8) | | |
| II | 2 (9.1) | | |
| III | 6 (27.3) | | |
| IV | 6 (27.3) | | |
| Unknown | 1 (4.5) | | |
| *Menstrual status (at time of sample collection), n (%)* | | | |
| Menstruating within past 3mo | 12 (54.5) | 11 (57.9) | 0.9802 |
| Natural menstrual cycles (within past 3mo) | 7 (31.8) | 7 (36.8) | |
| Hormone-induced cycles (within past 3mo) | 5 (22.7) | 3 (15.8) | |
| Amenorrheic (>3mo since last period) | 8 (36.4) | 6 (31.6) | |
| No response | 2 (9.1) | 2 (10.5) | |
| *Days since last menstrual period (at time of peripheral blood sample collection), n (%)* | | | |
| 0-7 days | 1 (4.5) | 0 (0.0) | 0.4708 |
| 8-14 days | 1 (4.5) | 0 (0.0) | |
| 15-35 days | 6 (27.3) | 5 (26.3) | |
| > 35 days | 3 (13.6) | 1 (5.3) | |
| Amenorrheic/not applicable | 8 (36.4) | 6 (31.6) | |
| No response | 3 (13.6) | 7 (36.8) | |
| *Days since last menstrual period (at time of EM surgical sample collection), n (%)* | | | |
| 0-7 days | 0 (0.0) | | |
| 8-14 days | 0 (0.0) | | |
| 15-35 days | 8 (36.4) | | |
| > 35 days | 3 (13.6) | | |
| Amenorrheic/not applicable | 8 (36.4) | | |
| No response | 3 (13.6) | | |
| *Reasons for hormone use, n (%)* | | | |
| Birth control | 12 (54.5) | 12 (63.2) | **0.0064*** |
| Pelvic pain or pain with periods | 15 (68.2) | 2 (10.5) | |
| Irregular periods | 6 (27.3) | 1 (5.3) | |
| Heavy periods | 11 (50.0) | 2 (10.5) | |
| Acne | 0 (0.0) | 2 (10.5) | |

| | | | |
|---|---|---|---|
| PMOS/PCOS | 3 (13.6) | 0 (0.0) | |
| Ovarian cyst | 9 (40.9) | 0 (0.0) | |
| Other | 6 (27.3) | 2 (10.5) | |
| No response | 2 (9.1) | 2 (10.5) | |
| *Hormones used for any reason (throughout lifetime), n (%)* | | | |
| Combined OCP | 10 (45.5) | 12 (63.2) | 0.0502 |
| Progestin only OCP | 2 (9.1) | 1 (5.3) | |
| OCP (unsure which one) | 8 (36.4) | 2 (10.5) | |
| Progestin injection | 3 (13.6) | 1 (5.3) | |
| Transdermal | 1 (4.5) | 1 (5.3) | |
| Vaginal ring | 2 (9.1) | 4 (21.0) | |
| Hormonal IUD | 9 (40.9) | 12 (63.2) | |
| Hormonal implant | 0 (0.0) | 0 (0.0) | |
| Oral progestins for cycle regulation | 12 (54.5) | 1 (5.3) | |
| GnRH agonist injection | 3 (13.6) | 0 (0.0) | |
| Norethindrone acetate | 1 (4.5) | 0 (0.0) | |
| Danazol | 1 (4.5) | 0 (0.0) | |
| Hormone replacement therapy | 1 (4.5) | 0 (0.0) | |
| Other | 2 (9.1) | 1 (5.3) | |
| Hormone (type unknown) | 4 (18.2) | 0 (0.0) | |
| None | 1 (4.5) | 0 (0.0) | |
| No response | 3 (13.6) | 1 (5.3) | |
| *Hormones used for any reason (within 3mo of sample collection), n (%)* | | | |
| Combined OCP | 1 (4.5) | 1 (5.3) | **0.0166*** |
| Progestin only OCP | 1 (4.5) | 0 (0.0) | |
| OCP (unsure which one) | 0 (0.0) | 0 (0.0) | |
| Progestin injection | 0 (0.0) | 0 (0.0) | |
| Transdermal | 0 (0.0) | 0 (0.0) | |
| Vaginal ring | 0 (0.0) | 0 (0.0) | |
| Hormonal IUD | 3 (13.6) | 10 (52.6) | |
| Hormonal implant | 0 (0.0) | 0 (0.0) | |
| Oral progestins for cycle regulation | 8 (36.4) | 1 (5.3) | |
| GnRH agonist injection | 1 (4.5) | 0 (0.0) | |
| Norethindrone acetate | 1 (4.5) | 0 (0.0) | |
| Danazol | 0 (0.0) | 0 (0.0) | |
| Hormone replacement therapy | 2 (9.1) | 0 (0.0) | |
| Other | 1 (4.5) | 0 (0.0) | |
| Hormone (type unknown) | 1 (4.5) | 0 (0.0) | |
| None (within 3mo) | 5 (22.7) | 7 (36.8) | |
| No response | 3 (13.6) | 1 (5.3) | |
| *Relevant prescription medications (ever taken for more than 3mo; throughout lifetime), n (%)* | | | |
| Thyroid drugs | 0 (0.0) | 1 (5.3) | 0.2578 |
| Drugs for RA | 0 (0.0) | 0 (0.0) | |
| Antibiotics (for 1mo or more) | 3 (13.6) | 0 (0.0) | |
| Antacids | 1 (4.5) | 2 (10.5) | |
| Drugs for stomach ulcer/gastritis | 4 (18.2) | 0 (0.0) | |
| Drugs for allergies (antihistamines) | 3 (13.6) | 2 (10.5) | |
| Steroids (oral, inhaled, or nasal) | 0 (0.0) | 1 (5.3) | |
| Inhaler for asthma | 1 (4.5) | 1 (5.3) | |
| None | 9 (40.9) | 11 (57.9) | |
| No response | 2 (9.1) | 3 (15.8) | |

| | | | |
|---|---|---|---|
| *Relevant prescription medications (ever taken for more than 3mo; taken daily at time of sample collection), n (%)* | | | |
| Thyroid drugs | 0 (0.0) | 1 (5.3) | 0.0709 |
| Drugs for RA | 0 (0.0) | 0 (0.0) | |
| Antibiotics (for 1mo or more) | 1 (4.5) | 0 (0.0) | |
| Antacids | 0 (0.0) | 0 (0.0) | |
| Drugs for stomach ulcer/gastritis | 2 (9.1) | 0 (0.0) | |
| Drugs for allergies (antihistamines) | 0 (0.0) | 0 (0.0) | |
| Steroids (oral, inhaled, or nasal) | 0 (0.0) | 0 (0.0) | |
| Inhaler for asthma | 0 (0.0) | 0 (0.0) | |
| None | 12 (54.5) | 13 (68.4) | |
| No response | 3 (13.6) | 3 (15.8) | |
| *Relevant OTC medications (used at least once a week for 3mo or longer; taken at time of sample collection), n (%)* | | | |
| Paracetamol/Acetaminophen | 11 (50.0) | 1 (5.3) | **<0.0001*** |
| Aspirin (325mg or more/tablet) | 1 (4.5) | 0 (0.0) | |
| Ibuprofen (e.g. Brufen) | 12 (54.5) | 1 (5.3) | |
| COX-2 inhibitors (e.g. Celebrex, Vioxx) | 0 (0.0) | 0 (0.0) | |
| Other anti-inflammatory analgesics (e.g. Naproxen, Mefenamic Acid, Aleve, Naprosyn, Relafen, Ketoprofen, Anaprox) | 4 (18.2) | 0 (0.0) | |
| Strong narcotic analgesics (e.g. Hydrocodone + Paracetamol, Codeine + Paracetamol, Morphine, Codeine, Oxycodone, Hydrocodone, Demerol) | 1 (4.5) | 0 (0.0) | |
| Other pain-killing drugs aimed at the nerves/CNS (e.g. Amitriptyline, Nortriptyline, Gabapentin, Pregabalin, Lamotrigine) | 3 (13.6) | (0.0) | |
| Muscle relaxants (e.g. Diazepam/Temazepam, Buscopan) | 2 (9.1) | 0 (0.0) | |
| Herbal Medicines | 2 (9.1) | 0 (0.0) | |
| None | 3 (13.6) | 14 (73.7) | |
| No response | 1 (4.5) | 0 (0.0) | |
| *Relevant comorbidity prevalence (throughout lifetime), n (%)* | | | |
| Asthma | 3 (13.6) | 1 (5.3) | 0.1705 |
| Crohn's Disease | 5 (22.7) | 3 (15.8) | |
| Graves' Disease | 3 (13.6) | 0 (0.0) | |
| Inflammatory Bowel Syndrome (IBS) | 0 (0.0) | 0 (0.0) | |
| Multiple Sclerosis | 1 (4.5) | 2 (10.5) | |
| Pelvic Inflammatory Disease (PID) | 0 (0.0) | 0 (0.0) | |
| PMOS/PCOS | 1 (4.5) | 0 (0.0) | |
| RA | 0 (0.0) | 0 (0.0) | |
| Sjögren's Syndrome | 0 (0.0) | 0 (0.0) | |
| Systemic Lupus Erythematosus (SLE) | 0 (0.0) | 0 (0.0) | |

| | | |
|---|---|---|
| Thyroid Disease | 0 (0.0) | 1 (5.3) |
| Ulcerative Colitis | 3 (13.6) | 1 (5.3) |
| None | 2 (9.1) | 6 (31.6) |
| No response | 2 (9.1) | 3 (15.8) |

CNS = central nervous system, GnRH = gonadotropin-releasing hormone, IUD = intrauterine device, OCP = oral contraceptive pill, OTC = over the counter, PCOS = polycystic ovary syndrome, PMOS = polyendocrine metabolic ovarian syndrome, RA = rheumatoid arthritis. *Statistically significant difference between EMS and control group.

**Table S2: Extracellular and intracellular antibodies used for flow cytometric analyses of PBMCs and PF cells.**

| Antibody | Fluorophore | Clone | Catalogue Number | Vendor |
|---|---|---|---|---|
| *PBMCs, Panel 1 – CytoFLEX S* | | | | |
| CD45 | Pacific Blue (PB) | HI30 | 304039 | BioLegend |
| CD4 | PE/Cy7 | RPA-T4 | 300511 | |
| CD196 (CCR6) | FITC | G034E3 | 353411 | |
| IL-17A | PE | BL168 | 512305 | |
| RORγt | APC | AFKJS-9 | 17-6988-82 | ThermoFisher |
| *PBMCs, Panel 2 – BD FACSAria III* | | | | |
| CD3 | APC | HIT3a | 300312 | BioLegend |
| CD4 | PE/Cy7 | RPA-T4 | 300511 | |
| IFNγ | FITC | N/A | 130-090-433 | Miltenyi Biotec |
| IL-17A | PE | N/A | 130-094-537 | |
| *PF cells, Panel 3 – BD FACSymphony* | | | | |
| CD45 | PE/Cy5.5 | HI30 | 35-0459-42 | ThermoFisher |
| CD4 | Brilliant Ultra Violet (BUV)™ 615 | RPA-T4 | 366-0049-41 | |
| CD8 | Qdot™ 655 | 3B5 | Q10055 | |
| CD25 | Super Bright (SB)™ 600 | BC96 | 63-0259-42 | |
| CD196 (CCR6) | PE-eFluor™ 610 | R6H1 | 61-1969-42 | |
| RORγt | APC | AFKJS-9 | 17-6988-82 | |
| IL-10 | BV711 | JES3-9D7 | 407-7108-41 | |
| FoxP3 | Brillant Violet (BV)™ 421 | 206D | 320123 | BioLegend |
| IL-17A | Alexa Fluor® (AF) 488 | BL168 | 512307 | |
| IFNγ | BV650™ | 4S.B3 | 502537 | |

**Table S3: FACS sorting of PBMCs and approximate cell yields sent for RNA sequencing (RNAseq).** CD4+ T cells magnetically isolated from PBMCs were sorted using FACS into respective cell populations: Th17 cells (CD3+CD4+IFNγ-IL-17+), Th1/17 (CD3+CD4+IFNγ+IL-17+), and Th1 cells (CD3+CD4+IFNγ+IL-17-).

| Sample ID | Total Thawed PBMC Cell Count | Total CD4+ T Cell Count (Inputted for FACS) | Total FACS Sorted Cell Counts (Sent for bulk RNAseq) |
|---|---|---|---|
| CON-030 | 5.30 x10$^6$ cells | 1.65 x10$^6$ cells | **Th17:** 7,909 cells |
| | | | **Th1/17:** 2,817 cells |
| | | | **Th1:** 93,539 cells |
| CON-031 | 6.35 x10$^6$ cells | 2.59 x10$^6$ cells | **Th17:** 22,635 cells |
| | | | **Th1/17:** 3,825 cells |
| | | | **Th1:** 113,194 cells |
| CON-032 | 6.45 x10$^6$ cells | 2.47 x10$^6$ cells | **Th17:** 12,143 cells |
| | | | **Th1/17:** 2,290 cells |
| | | | **Th1:** 74,422 cells |
| CON-033 | 18.15 x10$^6$ cells | 5.30 x10$^6$ cells | **Th17:** 25,355 cells |
| | | | **Th1/17:** 1,655 cells |
| | | | **Th1:** 78,749 cells |

| | | | |
|---|---|---|---|
| CON-034 | 11.85 $x10^6$ cells | 1.10 $x10^6$ cells | **Th17:** 6,167 cells |
| | | | **Th1/17:** 880 cells |
| | | | **Th1:** 50,163 cells |
| CON-035 | 12.80 $x10^6$ cells | 2.20 $x10^6$ cells | **Th17:** 17,117 cells |
| | | | **Th1/17:** 3,682 cells |
| | | | **Th1:** 116,154 cells |
| CON-036 | 11.80 $x10^6$ cells | 2.83 $x10^6$ cells | **Th17:** 59,613 cells |
| | | | **Th1/17:** 10,235 cells |
| | | | **Th1:** 242,812 cells |
| CON-037 | 6.60 $x10^6$ cells | 2.38 $x10^6$ cells | **Th17:** 24,348 cells |
| | | | **Th1/17:** 3,925 cells |
| | | | **Th1:** 123,508 cells |
| CON-038 | 7.15 $x10^6$ cells | 2.62 $x10^6$ cells | **Th17:** 25,987 cells |
| | | | **Th1/17:** 2,590 cells |
| | | | **Th1:** 126,443 cells |
| CON-039 | 8.10 $x10^6$ cells | 2.23 $x10^6$ cells | **Th17:** 9,661 cells |
| | | | **Th1/17:** 1,213 cells |
| | | | **Th1:** 62,807 cells |
| CON-040 | 17.35 $x10^6$ cells | 3.77 $x10^6$ cells | **Th17:** 14,508 cells |
| | | | **Th1/17:** 2,460 cells |
| | | | **Th1:** 122,392 cells |
| CON-045 | 12.85 $x10^6$ cells | 2.46 $x10^6$ cells | **Th17:** 10,498 cells |
| | | | **Th1/17:** 3,339 cells |
| | | | **Th1:** 154,473 cells |
| EMS-009 | 9.75 $x10^6$ cells | 2.41 $x10^6$ cells | **Th17:** 46,172 cells |
| | | | **Th1/17:** 8,016 cells |
| | | | **Th1:** 66,066 cells |
| EMS-014 | 11.00 $x10^6$ cells | 1.86 $x10^6$ cells | **Th17:** 12,483 cells |
| | | | **Th1/17:** 3,028 cells |
| | | | **Th1:** 68,375 cells |
| EMS-015 | 9.70 $x10^6$ cells | 1.70 $x10^6$ cells | **Th17:** 2,511 cells |
| | | | **Th1/17:** 732 cells |
| | | | **Th1:** 32,996 cells |
| EMS-016 | 2.49 $x10^6$ cells | 5.75 $x10^5$ cells | **Th17:** 6,636 cells |
| | | | **Th1/17:** 1,024 cells |
| | | | **Th1:** 23,408 cells |
| EMS-019 | 12.30 $x10^6$ cells | 3.22 $x10^6$ cells | **Th17:** 6,267 cells |
| | | | **Th1/17:** 2,072 cells |
| | | | **Th1:** 83,668 cells |
| EMS-022 | 14.10 $x10^6$ cells | 3.80 $x10^6$ cells | **Th17:** 40,048 cells |
| | | | **Th1/17:** 4,877 cells |
| | | | **Th1:** 97,054 cells |
| EMS-023 | 5.40 $x10^6$ cells | 1.52 $x10^6$ cells | **Th17:** 12,586 cells |
| | | | **Th1/17:** 7,398 cells |
| | | | **Th1:** 148,431 cells |
| EMS-024 | 7.80 $x10^6$ cells | 2.23 $x10^6$ cells | **Th17:** 33,381 cells |
| | | | **Th1/17:** 11,005 cells |
| | | | **Th1:** 99,013 cells |
| EMS-025 | 15.85 $x10^6$ cells | 3.53 $x10^6$ cells | **Th17:** 22,881 cells |
| | | | **Th1/17:** 5,638 cells |
| | | | **Th1:** 64,810 cells |
| EMS-026 | 6.10 $x10^6$ cells | 1.76 $x10^6$ cells | **Th17:** 6,232 cells |
| | | | **Th1/17:** 1,167 cells |
| | | | **Th1:** 58,976 cells |

CON = Control.
